# Identification of pathogenic variant enriched regions across genes and gene families

**DOI:** 10.1101/641043

**Authors:** Eduardo Pérez-Palma, Patrick May, Sumaiya Iqbal, Lisa-Marie Niestroj, Juanjiangmeng Du, Henrike Heyne, Jessica Castrillon, Anne O’Donnell-Luria, Peter Nürnberg, Aarno Palotie, Mark Daly, Dennis Lal

**Affiliations:** Cologne Center for Genomics, University of Cologne, Cologne, NRW, Germany; Luxembourg Centre for Systems Biomedicine, University Luxembourg, Esch-sur-Alzette, Luxembourg; Analytic and Translational Genetics Unit, Massachusetts General Hospital, Boston, MA, USA; Stanley Center for Psychiatric Research, Broad Institute of MIT and Harvard, Cambridge, MA, USA; Genomic Medicine Institute, Lerner Research Institute Cleveland Clinic, USA; Institute for Molecular Medicine Finland (FIMM), University of Helsinki, Helsinki, Finland; Epilepsy Center, Neurological Institute, Cleveland Clinic, Cleveland, USA

**Author notes:** **Correspondence: Dr. Dennis Lal**. **The authors declare no competing interests**.

## Abstract

Missense variant interpretation is challenging. Essential regions for protein function are conserved among gene family members, and genetic variants within these regions are potentially more likely to confer risk to disease. Here, we generated 2,871 gene family protein sequence alignments involving 9,990 genes and performed missense variant burden analyses to identify novel essential protein regions. We mapped 2,219,811 variants from the general population into these alignments and compared their distribution with 65,034 missense variants from patients. With this gene family approach, we identified 398 regions enriched for patient variants spanning 33,887 amino acids in 1,058 genes. As a comparison, testing the same genes individually we identified less patient variant enriched regions involving only 2,167 amino acids and 180 genes. Next, we selected *de novo* variants from 6,753 patients with neurodevelopmental disorders and 1,911 unaffected siblings, and observed a 5.56-fold enrichment of patient variants in our identified regions (95% C.I. =2.76-Inf, p-value = 6.66×10^−8^). Using an independent ClinVar variant set, we found missense variants inside the identified regions are 111-fold more likely to be classified as pathogenic in comparison to benign classification (OR = 111.48, 95% C.I = 68.09-195.58, p-value < 2.2e^−16^). All patient variant enriched regions identified (PERs) are available online through a user-friendly platform for interactive data mining, visualization and download at *http://per.broadinstitute.org*. In summary, our gene family burden analysis approach identified novel patient variant enriched regions in protein sequences. This annotation can empower variant interpretation.

## Introduction

Sequencing technologies are becoming routinely applied in clinical diagnostics(den Dunnen et al. 2016). The number of genetic variants derived from patients has increased exponentially(Lek et al. 2016), demanding scalable and accurate methods for variant interpretation. Particularly, the ability to accurately predict variants associated with rare and complex Mendelian disorders becomes crucial in the development of personalized medicine(Xue et al. 2015). Since 85% of disease traits are explained by variation within the coding portion of the genome, whole-exome and gene-panel sequencing have become the standard of care(Choi et al. 2009; Bamshad et al. 2011). Still, variant interpretation remains challenging(Gilissen et al. 2012), particularly for missense variants, which account for the most prevalent genomic alteration with 10,000 to 12,000 events per individual(Genomes Project et al. 2015). Protein truncating variants (PTVs) and large deletions are generally assumed to cause disease by loss-of-function mechanisms in haploinsufficient genes. In contrast, missense variants can have a variety of functional outcomes depending on the amino acid substitution and protein domain affected(Miosge et al. 2015) further complicating interpretation. Many computational tools have been developed for missense variant interpretation(Itan and Casanova 2015; Liu et al. 2016). These tools are based on a combination of criteria including the physicochemical properties of the amino acids in exchange (e.g., Grantham)(Grantham 1974), structural features (e.g., PolyPhen-2)(Adzhubei et al. 2010), amino acid conservation across different species (e.g., GERP++, SIFT)(Cooper et al. 2005; Kumar et al. 2009) or a combined machine learning consensus approaches (e.g., CADD, FatHMM, REVEL)(Shihab et al. 2013; Kircher et al. 2014; Ioannidis et al. 2016).

Repositories of variants from the general population have been used as a resource to calculate gene constraint or to identify coding regions “intolerant to variation”(Lek et al. 2016). Constraint metrics are extensively used for the identification of potential disease genes and for individual variant interpretation(Samocha et al. 2017). Missense variants are not randomly distributed across the exome, and functionally essential genes are constrained from variation(Petrovski et al. 2013; Bartha et al. 2018). Thus, variants within genes that are intolerant to loss of function or missense variation in the general population are more likely to be pathogenic(Lek et al. 2016). Methods and scores which incorporate variant tolerance have been developed. For example, the missense tolerance ratio (MTR) score evaluates constraint over a 31 amino-acid window(Traynelis et al. 2017); and the missense badness, polyphen-2, and constraint score (MPC) combines constraint with exchange and structural scores to report a missense specific score(Samocha et al. 2017).

From an evolutionary perspective, it has been shown that approximately 80% of Mendelian disorders have functionally redundant paralogs(Chen et al. 2013b) expressed across different cell types. Gene duplication events from ancestral genes have produced large sets of well-established paralog gene-families in the human genome, with different degrees of amino acid conservation and functional redundancy. Conserved amino acids across gene family members are more likely to hold essential functional domains. Thus, amino acid conservation across gene paralogs can be used at scale for variant interpretation (Parazscore)(Lal et al. 2017). Functional redundancy has led to the accumulation of pathogenic variants in analogous domains that are able to cause diseases in the tissues and organs the corresponding paralogous genes are expressed (Ware et al. 2012; Chen et al. 2013b; Walsh et al. 2014; Barshir et al. 2018). Therefore, protein alignments of gene family members could significantly cluster independent pathogenic variants in the same analogous domain. Variant aggregation over protein domain homologues without distinction of gene family members has been used for variant interpretation before(Gussow et al. 2016; Wiel et al. 2017), including the identification of cancer-driver variants(Melloni et al. 2016).

Similar to genetic constraint, patient variant clustering along the linear protein sequence can also be expected in functionally essential regions. Thousands of pathogenic variants have been used to train variant interpretation tools such as the variant effect predictor tool (VEST)(Carter et al. 2013) and the combined annotation dependent depletion (CADD)(Kircher et al. 2014). However, patient variant enrichment analysis to detect disease-sensitive regions has not been conducted on an exome-wide level.

Here, we compared the distribution of patient missense variants against population missense variants within gene family alignments and gene sequences alone. We developed a novel statistical framework that based on the observed mutational distribution, can identify pathogenic variant enriched regions (PERs) across protein sequences. We show that the family-wise approach is able to identify more and larger PERs than gene wise analyses. Our identified family-wise and gene-wise PERs are high in resolution and can be used for variant interpretation. We developed the “PER viewer” (http://per.broadinstitute.org) to facilitate the exploration of all data generated in this study, including gene family alignments, PERs, variants and paralog conservation scores in a user-friendly web application.

## Methods

### Population missense variants

Protein-coding variants from the general population were retrieved from the genome aggregation Database (gnomAD) public release 2.0.2(Lek et al. 2016). Exonic variants were downloaded in the Variant Call Format (VCFs) following gnomAD guidelines (http://gnomad.broadinstitute.org/downloads). Missense variants were extracted using vcftools(Danecek et al. 2011) based on the consequence “CSQ” field. The CSQ field is pre-annotated by gnomAD with the Variant Effect Predictor software (VEP, Ensembl v92) and provides information on 68 features, including gene/transcript, cross-database identifiers, as well as the desired molecular consequence. All annotations refer to the human reference genome version GRCh37.p13/hg19. Entries passing gnomAD standard quality controls (Filter = “PASS” flag) and annotated to a canonical gene transcript (CSQ canonical = “YES” flag) were extracted. The canonical transcript is defined as the longest CCDS translation with no stop codons according to Ensembl(Zerbino et al. 2018). Missense variants calls were merged into one single file matching genomic position and annotation. The final “Population” dataset contains all missense variants within canonical transcripts found in the general population.

### Patient missense variants

Disease-associated missense variants were retrieved from two sources: The ClinVar database (ClinVar), December, 2017 release(Landrum et al. 2016), and The Human Gene Mutation Database (HGMD®) Professional release 2017.2(Stenson et al. 2003). ClinVar variants were downloaded directly from the ftp site (ftp://ftp.ncbi.nlm.nih.gov/pub/clinvar/) in a table format. Molecular consequence was inferred through the analysis of the Human Genome Variation Society (HGVS) sequence variant nomenclature field(den Dunnen et al. 2016). Specifically, when the variant was reported to cause an amino acid change different to the reference, it was subsequently annotated as a missense variant (e.g. *p.Gly1046Arg*). To increase stringency, only “Pathogenic” and/or “Likely Pathogenic” classified ClinVar variants were kept. The HGMD data was directly filtered for “missense variants”, “High Confidence” calls (hgmd_confidence = “HIGH” flag) and “Disease causing” state (hgmd_variantType = “DM” flag). All annotations refer to the human reference genome version GRCh37.p13/hg19, and variants belonging to non-canonical transcripts were removed. Since ClinVar and HGMD are not mutually exclusive, we took the union of both resources and removed duplicated entries by comparing HGVS annotations. The final “Patient” dataset contains patient derived missense variants and their corresponding disease annotation.

### Gene-family definition

Gene families were retrieved following the method described in Lal et al, 2017(Lal et al. 2017). Briefly, we downloaded the human paralog definitions from the Ensembl BioMart system(Kinsella et al. 2011). Noncoding genes and genes without an HGNC(Yates et al. 2017) symbol were excluded. Similarly, gene families with less than two HGNC genes were filtered out. For all analyses, we used one transcript per gene, keeping only the canonical version according to Ensembl. To construct a family-wise FASTA file, for all canonical transcripts respective consensus coding sequences (CCDS) were downloaded from the UCSC table browser(Karolchik et al. 2004). Family protein sequence alignment was conducted with MUSCLE(Edgar 2004). Younger evolutionary paralogs show higher functional redundancy(Chen et al. 2013a). To avoid alignments of strongly diverging sequences and to enrich for overall similarity, we filtered out families with less than 80% similarity in their overall protein sequence(Dufayard et al. 2005). In total, we used 2,871 gene families comprising 9,990 genes. Family names and gene members are available at our GitHub repository (https://github.com/dlal-group/per).

### Missense Variant mapping

The population and patient missense variants were independently mapped to corresponding amino acids in all gene family protein sequence alignments. Population and patient missense variant mapping was conducted using a binary annotation: “0” for amino acids with no missense variant reported, and “1” for residues with at least one missense variant reported. We expected that constrained regions across the gene family alignment will be enriched with amino acids marked as “0”, whereas, disease-sensitive regions will cluster amino acids marked with “1”. Gene family alignment regions with more gaps are less conserved than aligned amino acids and are more likely to be not functionally essential. Thus, in the population variant mapping, gaps introduced in any gene family member were also assigned a “1” state as if they were mutated to punish less conserved sites. For the patient variant mapping the gaps were kept as “0”. Since every missense variant contained in the patient subset was associated with at least one phenotype in one gene, multiple genes and diseases were aggregated in aligned residues upon alignment. This information was collected in an additional “*Gene:Disease*” field for further follow up. Population and patient datasets containing all missense variants analyzed in the present study as well as the Perl scripts used to filter and finally map the variants in to the alignments are available at our GitHub repository (https://github.com/dlal-group/per).

### Missense burden analysis – gene family alignments

We performed statistical comparisons between population and patient variants mapped to protein family alignments. Specifically, we applied sliding windows of 9 amino acids across index positions of the paralog alignments with a 50% overlap to increase sensitivity (Figure 1a). We summed the number of “0” and “1” sites inside and outside the window across the whole alignment index. A one-sided Fisher’s exact test with 95% confidence was performed over each sliding window, comparing general population and patient counts inside the window against the corresponding counts outside of it. For example, a burden analysis based on a sliding window of size 5 will first test the counts of index positions 1 to 5 against the counts found from position 6 to the end of the alignment (Figure 1). Bonferroni multiple testing adjustment was applied accounting for the total number of sliding windows tested for each gene family alignment. Sliding windows with adjusted p-values below 0.05 were considered significant and subsequently called *Pathogenic Enriched Regions* (PERs). If two or more consecutive sliding-windows were found significant, the final PER reported will reflect the fusion of all consecutive significant windows boundaries. In order to identify the optimal sliding window size, the analysis was executed with multiple sliding window sizes, from 3 up to 31 amino acids, to evaluate the window size sensitivity and specificity (**Supplementary Figure 1**). Sensitivity was measured by the number of significant regions detected, amino acids involved, and gene-families affected. Specificity of the analysis was measured by the ratio between the number of amino acids sites inside PERs with no disease associations, and the number of amino acids inside PERs with disease associations (i.e., in at least one family gene-member). Missense variant mapping and sliding window counts were performed with an in-house Perl script. Fisher’s exact tests, Bonferroni adjustment and plots were performed with the R statistical software(R Core Team 2016).

**Figure 1.**
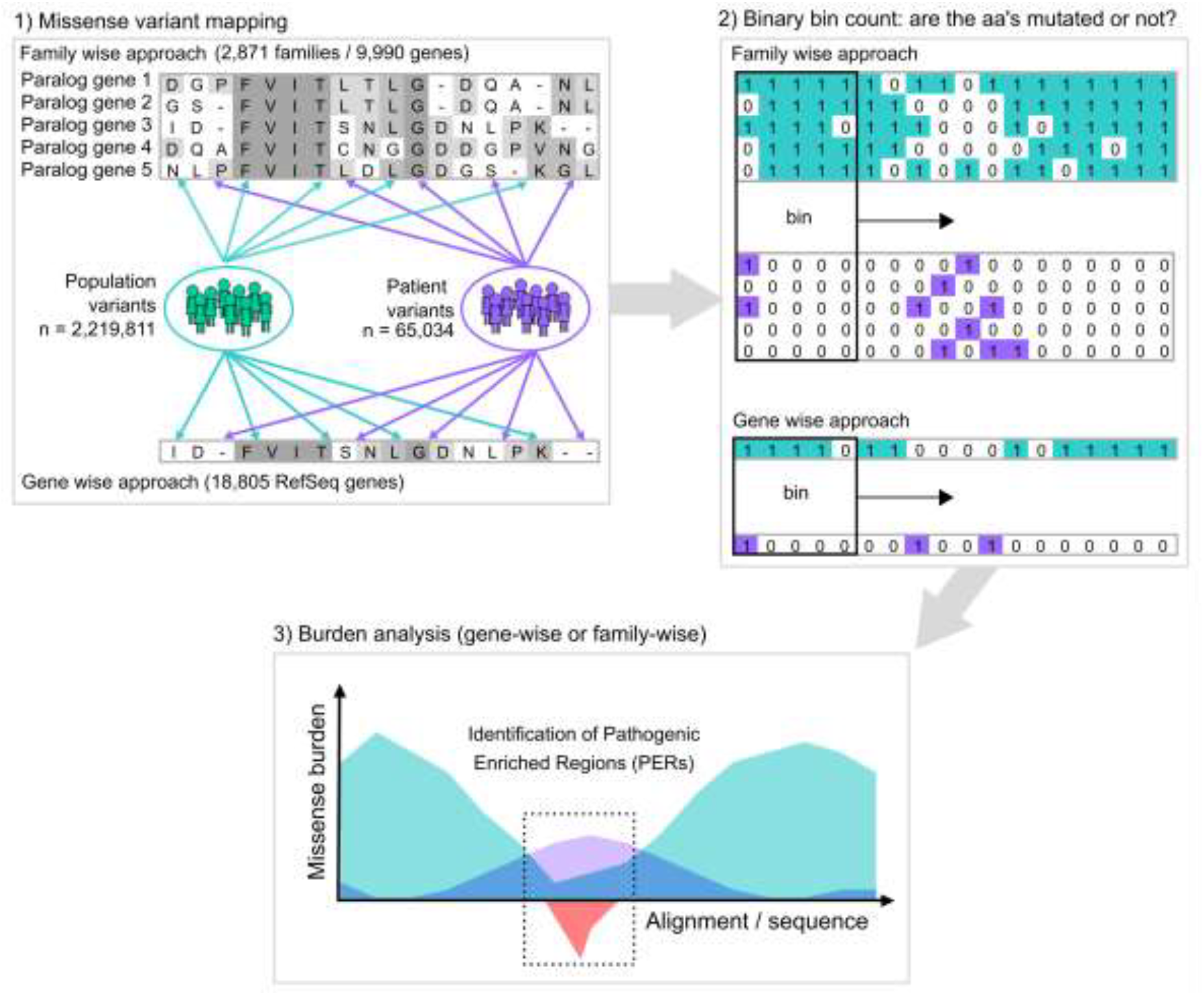
Study workflow and the PER viewer. **a.** Study workflow: starting from protein alignments of paralogs genes (gene family approach) or all genes (gene-wise approach) missense variants from gnomAD (population, green) and ClinVar/HGMD (patient, purple) were mapped independently to the corresponding amino acid positions (binary notation). For sites with at least one missense variant reported a “1” state was assigned. Alternatively, if no mutation was found a “0” state was annotated instead. Amino acid sliding window counting over the alignment/sequence was used to calculate the corresponding missense burden. Statistical comparison between the neural-control and patient burden (green area vs purple area) allowed the identification of significant regions in both approaches (red area).

### Missense burden analysis – Gene-wise analysis

The *Missense variant mapping* and *Burden analysis* protocols were further applied to all RefSeq genes independently to evaluate gene-wise enrichment. For all 18,806 canonical transcripts, their respective CCDS was downloaded from the UCSC table browser(Karolchik et al. 2004). The missense variant mapping and burden analyses were conducted using the same Perl scripts, treating each gene as a one-member “family”. Gene names, identifiers, and corresponding canonical protein sequences are available at our GitHub repository (https://github.com/dlal-group/per).

### Development of PER viewer

Population and patient missense burden calculations as well as the identification of significant regions within genes and gene families were made publicly available through the PER viewer (*http://per.broadinstitute.org*). PER viewer was developed with the Shiny framework of R studio(Team 2015), which transforms regular R code into HTML that can be displayed by any web browser. Pre-calculated burden analyses for all genes and gene-families were deployed in a Google Virtual Machine (VM) using the googleComputeEngineR(Edmondson 2017) package. All graphs shown in the present work and by the online tool are based on the ggplot2 R library(H. 2009).

## Results

### Missense variant mapping in genes and gene-families

Protein residues near or within clusters of pathogenic variants are more likely to be disease associated. Our goal was to generate an annotation for protein regions vulnerable to disease. We compared the distribution of missense variants from patients and individuals from the general population across protein sequences to identify regions enriched for patient variants. To increase our statistical power, we performed a ‘family-wise’ approach analyzing the missense variant burden along aligned protein sequences of gene-family members (i.e. paralogs) as a single unit. First, we extracted a total of 2,219,811 missense variants from the Genome Aggregation Database(Lek et al. 2016) (gnomAD) to serve as our ‘Population’ variant dataset. Patient missense variants were retrieved from two sources: the ClinVar database(Landrum et al. 2016) and the Human Gene Mutation Database (HGMD)(Stenson et al. 2003). After variant filtering, the union of ClinVar and HGMD yielded a total of 65,034 unique high-confidence pathogenic/likely-pathogenic missense variants which were subsequently used as our “Patient” dataset. A detailed description of the applied filtering criteria can be found in the Methods section.

### Missense burden analysis

To investigate mutational burden across paralog conserved amino acids, we mapped all missense variants from the Population and Patient datasets onto a set of 2,871 gene-family protein alignments involving 9,990 genes. To generate a “gene-wise” analysis as a comparison group, we applied the same mapping procedure to the single protein sequences of 18,805 RefSeq genes. The workflow designed for PER detection is summarized in Figure 1. To calibrate the optimal sliding window size, we conducted multiple rounds of burden analyses with varying window sizes (see Methods). We observed that at greater sliding window sizes, more PERs were detected; however, the ratio of aligned amino acids positions with patient variants versus without was decreasing (**Supplementary Figure 1**).

To ensure specificity, we decided to limit PERs to contain a minimum of 50% of amino acids with at least one disease association in any gene family member. As a result, the analysis was calibrated to a sliding window of nine amino acids (**Supplementary Figure 1c**).

### PER detected in the family-wise and gene-wise approaches

We identified 398 and 210 PERs in the family-wise and gene-wise analysis encompassing 33,887 and 2,167 amino acids, respectively (Figure 2a). Collectively, a total of 34,890 amino acids from 1,126 genes fall within PERs boundaries which can be traced back to 104,670 nucleotides in the reference genome. Genomic coordinates and annotation of all identified PERs are available at our github site (https://github.com/dlal-group/per). We observe a 5.8-fold enrichment of genes with at least one PERs in the family wise analysis (n = 1,058) than in the gene-wise analysis (n = 180). All genes in the gene-wise approach have been previously associated to disease, however, the family-wise approach was able to detect PERs in 560 genes not yet associated with a human phenotype (Figure2b). Similarly, among the amino acid positions within family-wise PERs, 86.2% (n=29,222) have no prior disease association in comparison to 53.5% (n=1,161) of the gene wise PERs (Figure 2c). In general, the family-wise approach identified more PERs, more amino acids sites and patient variants than the gene-wise approach. Overall, 84% of genes with at least one PER were identified exclusively through the family-wise approach, while 6.2% of the total genes (n = 70) were found exclusively in the gene-wise analysis. A total of 110 genes (9.8%) had PERs in both approaches (Figure 2d). It is important to note that 64 out of the 70 (91.4%) genes with PERs exclusively found in the gene-wise analysis do not belong to a gene family. The average patient variant fold enrichment observed for PERs identified in the family-wise approach was lower than that observed for PERs identified by the gene-wise analysis (Figure 2e). However, the corresponding association was more significant in the family-wise analysis compared to the gene-wise analysis (Figure 2f).

**Figure 2.**
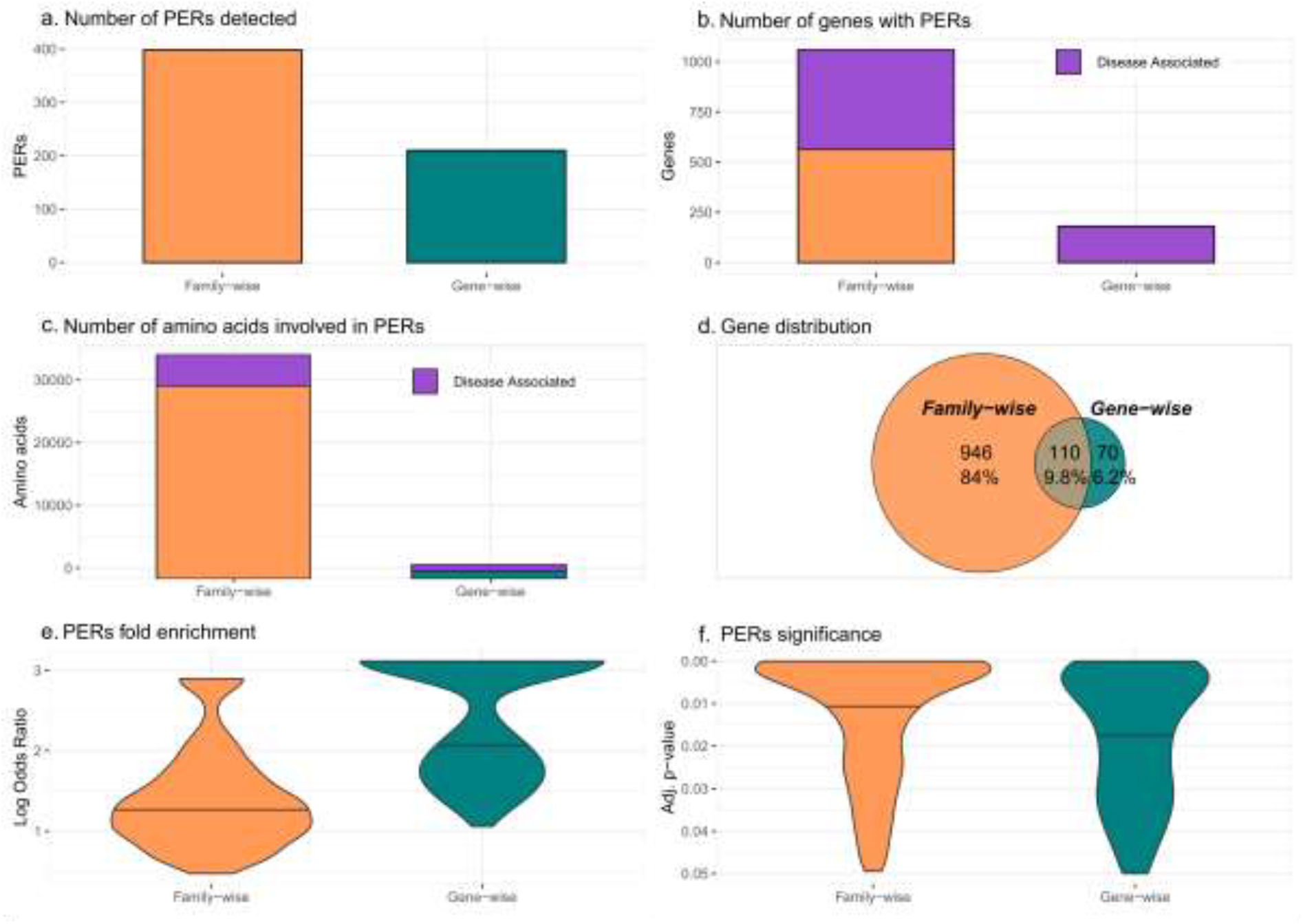
PERs detected with the family-wise and gene-wise burden analyses. Summary statistics for family-wise (orange) and gene-wise (green) approaches are shown for: **a.** Number of PERs detected, **b.** Number of genes with PERs and **c.** Number of amino acids involved in in PERs. For b and c the number of genes and amino acids associated to disease is shown in purple. Next, in **d.** to reflect gene distribution a Venn diagram is shown with the proportion of genes with PERs exclusively found in both approaches and the ones that are shared between them. Overall enrichment (Log Odds Ratio) and significance (adjusted p-value) distribution of all PERs detected in each approach are shown in **e**. and **f**., respectively.

### Illustrative Example: The voltage-gated sodium channel gene family

We show the missense burden analysis results of the voltage-gated sodium channel gene family (family ID: 2614) composed of ten paralogous genes: SCN1A, SCN2A, SCN3A, SCN4A, SCN5A, SCN7A, SCN8A, SCN9A, SCN10A, and SCN11A (Figure 3). The alignment of the ten protein sequences consists of 2,188 amino acids alignment index positions, onto the patient and population variants were subsequently mapped. Clinical phenotypes from patient variants found in any gene-family member were aggregated into the corresponding aligned amino acid position. The missense burden analysis identified 15 PERs in the Voltage-Gated Sodium Channel Family (Figure 3a). Overall, regions with a drop in the distribution of population variants are increased for patient variants and vice versa. PER9 represented the longest patient variant enriched region, with 29 aligned amino acids sites from positions 1,476 to 1,504. As an example, we show the clinical phenotypes of patients carrying variants in PER2, located between aligned positions 456 to 464 (Figure 3b), which can be explored using the table browser feature (Gene:Disease column). The patient variant enrichment within PER2 was based on missense variants in SCN5A (n=5), SCN4A (n=4), SCN1A (n=4), SCN8A (n=3) and SCN2A (n=2) representing patients with Long QT syndrome, Brugada syndrome, Dravet syndrome, and a broad range of infantile epilepsies and epileptic encephalopathies. In contrast to the family-wise burden analysis, the gene-wise burden analysis was not able to identify PERs in any of the ten voltage-gated sodium channel genes (**Supplementary Figure 3**), indicating greater statistical power of family-wise burden analysis in this gene family.

**Figure 3.**
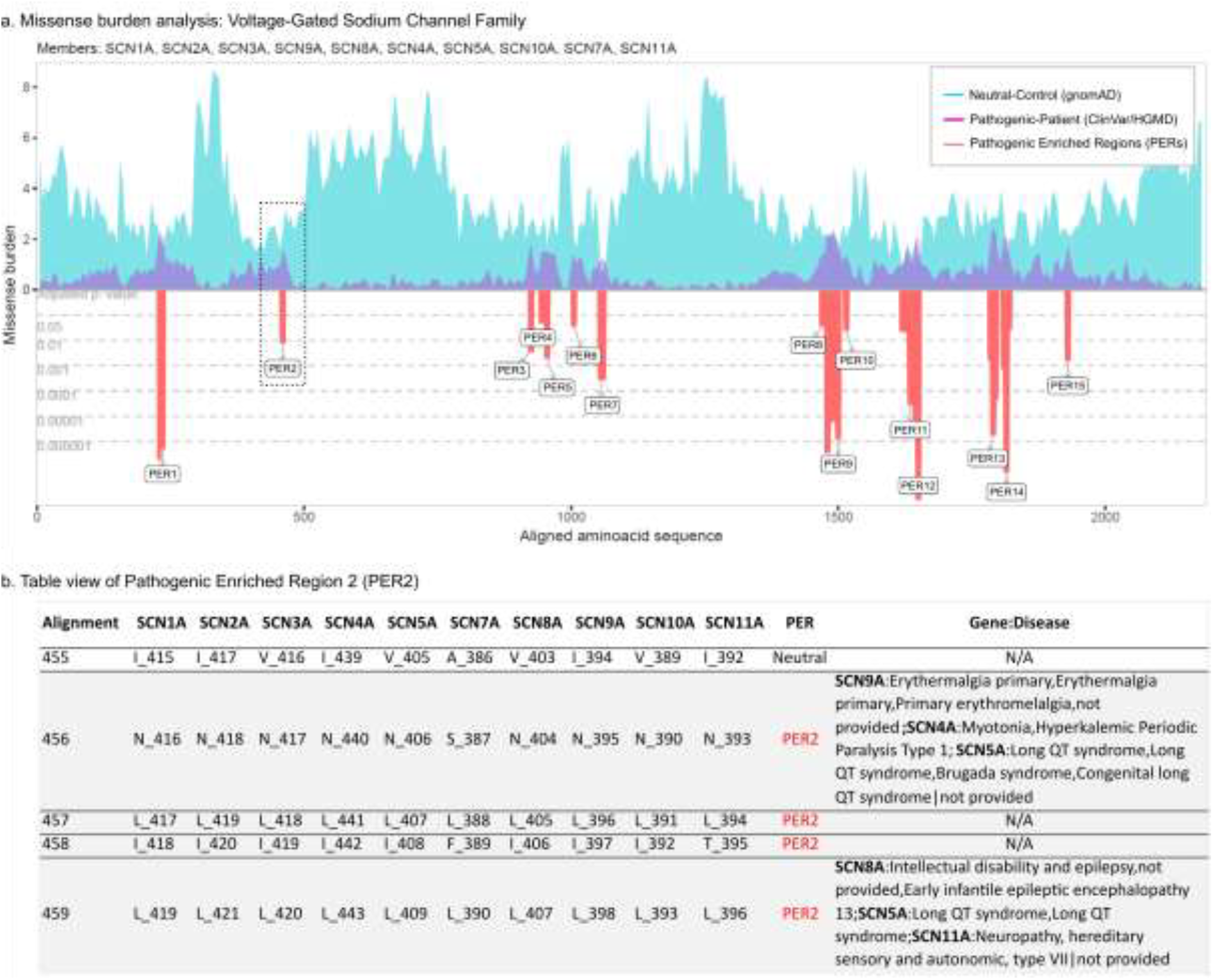
PER viewer tool example: The Voltage-Gated Sodium Channel Family. **a.** Missense burden analysis of the voltage-gated sodium channel protein family (Family ID: 2,614.subset.3) composed by *SCN1A*, *SCN2A*, *SCN3A*, *SCN4A*, *SCN5A*, *SCN7A*, *SCN8A*, *SCN9A*, *SCN10A*, *SCN11A*. Population and patient missense burden are shown in green and purple, respectively. Significant pathogenic enriched regions identified (PERs) are shown in red negative area and proportional to their adjusted p-values (gray horizontal lines). **b.** Table view of pathogenic enriched region 2 (PER2, Positions 455-459). Gene columns denote individual canonical sequence alongside corresponding amino acid position. Column “Gene:Disease” displays analogous diseases observed in the patient dataset. N/A sites shows aligned amino acids positions with no disease reported.

### PER viewer

We developed an R-based online tool to make the full set of results accessible: “PER viewer*”* available at http://per.broadinstitute.org. The main features of PER viewer are shown in **Supplementary Figure 2**. The user can query any gene and search for its corresponding missense burden analysis results. If the gene belongs to a gene-family, the results will be shown family-wise with the option to evaluate genes independently.

For genes which do not belong to a gene-family, the single gene burden analysis will be shown instead. Burden analyses and table browsing are displayed in the same format shown for the sodium channel family (Figure 3). The user can explore the burden observed in the Population and Patient datasets along the alignment or gene sequence at an amino acid level. Alignments, burden analyses, summary statistics, and the identification of PERs are available for download at PER viewer.

### PERs on independent cohorts

To test the utility of PER annotation in an independent dataset, we evaluated the distribution of *de novo* missense variants (DNVs) within and outside of the identified PERs from a large neurodevelopmental (NDD) case control cohort(Heyne et al. 2018). The dataset included 6,753 patients with 4,404 missense DNVs identified and 1,911 unaffected siblings with 768 missense DNVs identified (Figure 4a). Patient missense DNVs (n=185) were 5.56-fold enriched within PERs compared to control missense DNVs (OR= 5.56, 95% C.I. =2.76-Inf, p-value = 6.66×10^−8^). The fold enrichment of patient variants in PERs was even greater when we restricted the analysis to constrained genes (pLI >0.9)(Lek et al. 2016). For this group of haploinsufficient genes, a 17.98-fold patient DNV enrichment was observed (OR=17.98, 95%C.I. = 3.70-Inf, p-value = 1.29 x10^−6^). Notably, it is not expected that all patient DNVs are pathogenic. In an additional analysis, we evaluated the distribution of missense variants reported in later ClinVar release (January, 2019). We found 4,605 ClinVar variants inside PERs boundaries. While 43.88% of ClinVar missense variants inside PERs are classified as pathogenic (n = 2,021), only 16 benign missense variants are found within the boundaries of PERs (Figure 4b, **orange bars**). We compared the number of ClinVar pathogenic and benign variants inside and outside PERs, we observed a 111-fold enrichment for pathogenic variants (OR= 111.14, 95%C.I = 68.09-195.58, p-value < 2.2e-16). We also observed 1,082 missense variants within PERs that at currently classified as variants of unknown significance (VUS).

**Figure 4.**
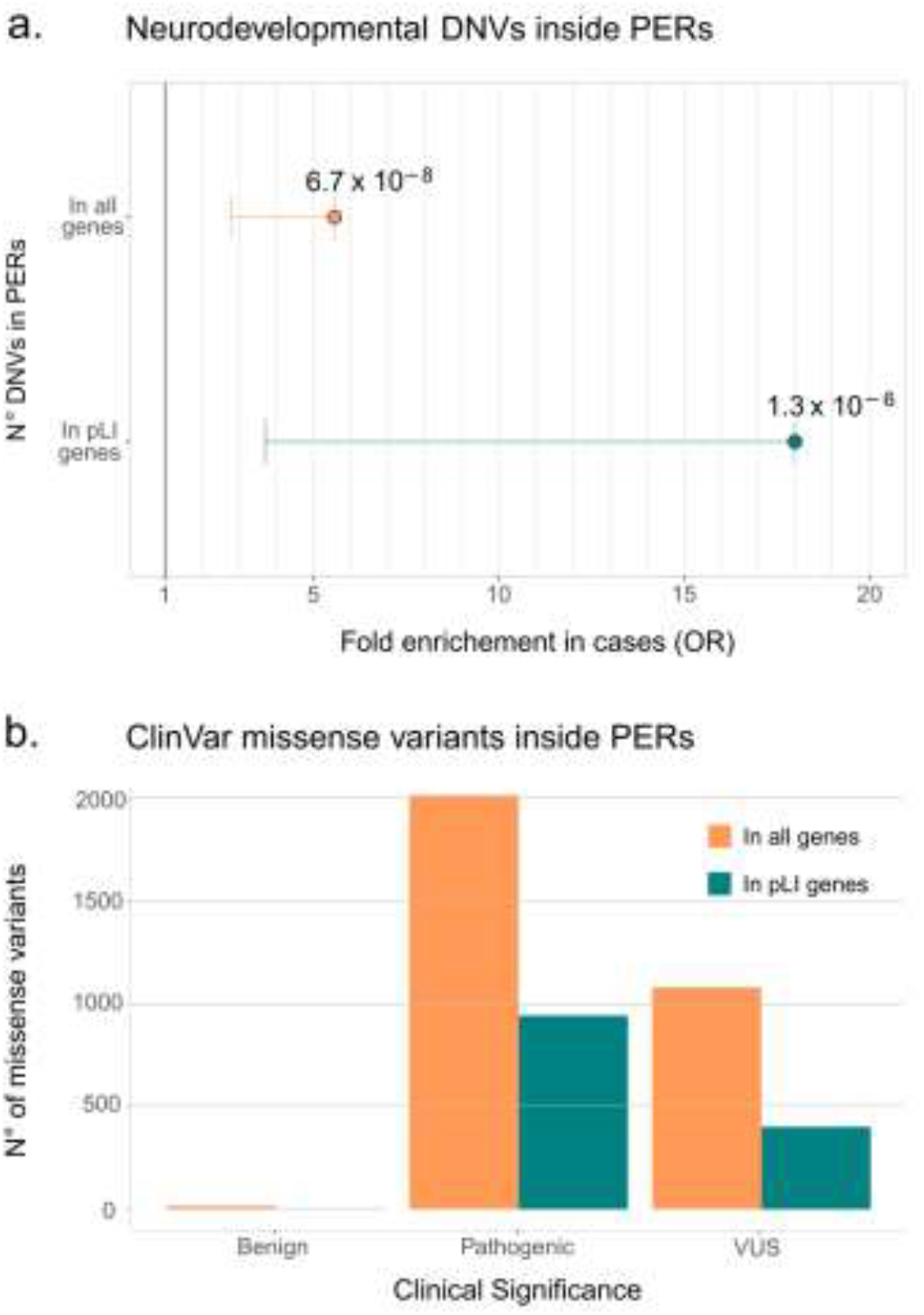
Disease-causing variants are enriched in PERs. **a.** Neurodevelopmental Disorder DNVs inside PERs. Case and Control comparison of *de novo* variants (DNVs) inside PERs retrieved from Heyne et al, 2018(Heyne et al. 2018) is shown for all genes (orange, OR= 5.56, 95% C.I. =2.76-Inf, p-value = 6.66×10^−8^) and genes with high probability of being loss-of-function intolerant (pLI >0.9, green, OR=17.98, 95%C.I. = 3.70-Inf, p-value = 1.29 x10-6). Fold enrichment observed in cases was calculated with a one-sided Fisher’s exact test. Resulting odds ratio (OR) with 95% confidence and corresponding p-values are shown in the horizontal axis. **b.** ClinVar missense variants (from Jan, 2019 release) inside PERs with Benign, Pathogenic and Unknown (VUS) clinical significance. Number of variants observed is shown considering all genes (orange) and pLI >0.9 genes only (green).

## Discussion

The present work compares the exome-wide distribution of missense variants from the general population with patient variants across single protein sequences and gene family protein sequence alignments. The family-wise approach was more sensitive and powerful than the basic gene-approach in PERs detection (Figure 2). We show that missense variants in amino acid positions within PERs are more likely to be classified as pathogenic rather than benign. These regions, enriched for patient variants and depleted for population variants, likely encompass functionally essential protein features. Thus, the generated exome-wide map of PERs can be used as an additional criterion for variant interpretation. Specifically, PER annotation and evaluation could be included in the “PM1” category of the American College of Medical Genetics (ACMG) guidelines. PM1 is defined as “variants located in a mutational hot spot and/or critical and well-established functional domain without benign variation”(Richards et al. 2015). Furthermore, the statistical framework designed to detect PERs provides fold enrichments and 95% confidence intervals that can be integrated in to Bayesian tools based on ACMG guidelines(Tavtigian et al. 2018). It has been estimated that an observed fold enrichment above 18.7 can be considered as a strong criteria for variant interpretation. Thus, 31.81% of all PER sites (33,300 nucleotides) could be further incorporated as a strong criteria for variant interpretation.

Identification of functional essential domains and sites across single protein sequences represents a challenge for rare Mendelian disorders. The number of patient variants annotated for most genes is still small and limits variant interpretation and prediction score development. Our approach aggregates variants across analogous sites within gene families to single unit hypothesizing that functionally essential sites across related proteins are conserved. We observed that the distribution pattern across protein sequences of patient and population variants was similar across gene family members, which yielded in a larger number of PERs and genes with PERs in the family approach compared to the single gene approach (Figure 2). Similar sequence grouping approaches have been conducted over homologous protein domains(Wiel et al. 2017), defined as functional sub-units that can be present in a broad spectrum of unrelated proteins(Finn et al. 2016). In contrast to domain-wise approaches, our missense burden analyses was performed on functionally redundant genes. Paralogous genes have accumulated a significant amount of disease variants because they can be masked by paralog functional redundancy(Chen et al. 2013b; Barshir et al. 2018). Paralog families can leverage additional insights in the context of sequence grouping approaches.

In contrast to other variant interpretation tools such as MTR(Traynelis et al. 2017), VEST 3.0(Carter et al. 2013) or CADD(Kircher et al. 2014), PER viewer does not provide a score for all possible missense mutations, but instead a set of amino acids regions where pathogenic variants accumulate significantly. PERs are able to capture aligned amino acids sites, irrespective of disease association, which allows variant interpretation even if no missense variant has been previously reported (Figure 2b, e.g. alignment index position 457). The family-wise variant annotation allows to manually inspect variants across the alignment index position, which can be useful for biological and clinical interpretation of variants. For example, in the voltage-gated sodium channel gene family, we observe at the filly paralog conserved PER2 index position 456 (Figure 2b) pathogenic classified variants for genes *SCN9A*, *SCN4A* and *SCN5A.* Thus, potential variants at the same alignment index position found in the future in the genes *SCN8A or SCN11A* (Asparagine 404 or asparagine 393, respectively) are more likely to be pathogenic, reflecting the practical use of our family-wise approach. In fact, paralogous annotation and variant interpretation transfer has been explored before and could be considered as common practice in the electrophysiology field(Ware et al. 2012; Walsh et al. 2014).

Our approach has several limitations. The statistical identification of PERs is dependent on the number of identified variants. We cannot rule out if missense variants outside PER boundaries are pathogenic, rather we are prioritizing variants within these regions. It is likely that more PERs will be identified as the power of this approach increased as population and patient databases continue to grow. Furthermore, paralogs belonging to the same family may evolve different functions through the development of specific domains(Pires-daSilva and Sommer 2003; Dos Santos and Siltberg-Liberles 2016). Upon alignment, gene specific domains will not show conservation, thereby having less likelihood to reach significance in the family-wise burden analysis. Nevertheless, if gene specific domains are in fact enriched for pathogenic variants, the gene-wise approach could still identify PERs in such regions. Moreover, functional redundancy among paralogs, does not guarantee the same degree of tolerance or intolerance to variation. Burden analyses including genes tolerant to variation will introduce noise and may mask specific signals. Additional filtering of gene family members may improve specificity in larger gene families, losing smaller gene families in the process. Our approach is able to identify protein regions constrained for variants in the general population and likely disease causing when mutated. Protein regions which can confer risk to disease through low penetrance variants or late onset of disease after typical reproductive age are unlikely to be identified due to little constraint in the general population(Bodmer and Bonilla 2008). The missense burden analysis is highly dependent on the number and quality of variants used as reference for population and patient dataset. We cannot rule out that the ascertainment of variants for specific genes can be skewed - e.g., different sequencing coverage of patient and population variants. However, further releases of gnomAD and ClinVar/HGMD will add even more power to the analysis. Notably, our framework is not restricted to the aforementioned resources and can be implemented with alternatives inputs. For example, missense burden analysis using the catalogue of somatic mutations in cancer (COSMIC)(Forbes et al. 2017) may be of use in the detection of cancer specific PERs.

The American college of medical genetics guidelines (ACMG)(Richards et al. 2015) has made considerable efforts to provide guidelines and standardize criteria for pathogenicity assignment. Nevertheless, about 60.8% of missense variants in ClinVar (release January 2019) have either conflicting reports of pathogenicity, no interpretation at all or are annotated as variants of unknown significance (VUS)(Landrum et al. 2016). With increasing data, machine learning approaches are likely to outperform older variant prediction algorithms such as PolyPhen and SIFT. Recently, studies that show this trend have been published(Itan and Casanova 2015). However, they lack the ability to understand why a given prediction score is high or low - limiting translation into therapeutics and biology. With the PER viewer, we are able to collect the phenotypes observed on a given region in an online tool that can simultaneously serve as a intuitive variant interpretation tool. PERs will empower gene discovery studies by facilitating the identification of specific regions within these candidate disease genes. This will have immediate impact on prioritizing candidate variants researchers and molecular diagnostic labs that will evaluate variants within PERs and genes with PERs more carefully.

## Acknowledgments

We thank the Genome Aggregation Database (gnomAD) and the groups that provided exome- and genome-wide data to this resource. A full list of contributing groups can be found at http://gnomad.broadinstitute.org/about. We also thank the researchers and clinicians involved in the generation of clinical data contained in ClinVar and HGMD as well as the respectives patients and their families. EP was supported by Dravet Foundation research grant to DL.

## Author Contributions

EP-P, and DL conceived and designed the study; PM, AP, AO-L, PN and MD supervised the study; EP-P, SI, LMN, JC and HH gathered and analyzed the data; EP-P and JD designed and developed the web interface; EP-P, PM and DL drafted the manuscript with input from all authors.

